# Transcriptionally defined morphological subtypes of pancreatic ductal adenocarcinoma

**DOI:** 10.1101/2022.09.23.509133

**Authors:** Teresa G Krieger, Alexander Sudy, Felix Schicktanz, Luca Tosti, Johannes Liebig, Björn Konukiewitz, Morgane Rouault, Anežka Niesnerová, Xiaoyan Qian, Wilko Weichert, Roland Eils, Katja Steiger, Christian Conrad

## Abstract

Tumour heterogeneity remains a major obstacle to effective and precise therapy for pancreatic ductal adenocarcinoma (PDAC), the most common pancreatic cancer. Several transcriptional subtypes of PDAC with differential prognosis have been described, but they co-occur within tumours and are difficult to distinguish in routine clinical workflows. To investigate the relationship between transcriptional PDAC subtypes, local tissue morphology and the tumour microenvironment, we employed in situ sequencing to profile single cells in their spatial tissue context. We identify five transcriptional subtypes of PDAC cells occurring in three distinct morphological patterns, including secretory tumour cell monolayers, invasive tumour cells with high expression of cell adhesion molecules *CEACAM5* and *CEACAM6*, and spatially distributed tumour cells associated with inflammatory-type fibroblasts. Analysis of bulk RNA-sequencing datasets of the TCGA-PAAD and PACA-AU cohorts according to these spatio-transcriptional subtypes confirmed their prognostic significance. Our results thus indicate an automatable substratification based on spatially-resolved transcriptomics of PDAC and identify distinct subtypes of ‘classical’ PDAC, representing most cases of this devastating malignancy.

## Introduction

Pancreatic ductal adenocarcinoma (PDAC) is the most lethal of all major organ malignancies, with a 5-year survival rate of less than 10% ^1,2^. Due to a lack of treatment advances compared to other cancer types, PDAC is predicted to become the second leading cause of cancer deaths in the United States by 2030 ^3^.

Bulk transcriptomic analyses have converged on two transcriptional subtypes of PDAC tumours, termed ‘classical’ and ‘basal-like’, with prognostic significance as the latter carry a poorer prognosis ^4^. Related molecular subtyping schemes have also been suggested, including a distinction into ‘classical’, ‘quasi-mesenchymal’ and ‘exocrine-like’ PDAC ^5^ or into ‘squamous’, ‘pancreatic progenitor’, ‘immunogenic’ and ‘aberrantly differentiated endocrine (ADEX)’ tumours ^6^. These schemes show significant overlap as the ‘classical’ and ‘pancreatic progenitor’ subtypes, as well as the ‘basal-like’, ‘quasi-mesenchymal’ and ‘squamous’ subtypes, share similar transcriptional signatures ^7^. Due to low neoplastic cellularity in tumour samples, a recent study reported that the ‘exocrine-like’, ‘immunogenic’ and ‘ADEX’ subtypes may represent contaminating non-neoplastic cells instead of tumour cells ^8^.

Recent single-cell transcriptomics studies provide emerging evidence that molecular PDAC subtypes are not mutually exclusive, but co-occur within the same tumours ^9,10^. Instead of discrete tumour cell states, PDAC cells may thus occupy a continuum of tumour cell states ranging from ‘classical’ to ‘basal-like’, requiring further investigation ^11^.

Histopathologically, PDAC tumours are graded according to defined WHO criteria that include the occurrence of tubular duct-like structures or solid areas, retained mucin, nuclear polymorphism and number of mitoses ^12^. A recent histopathological investigation distinguished ‘gland forming’ and ‘non-gland forming’ components based on the presence or absence of well-formed glands, and showed that tumours with at least 40% ‘non-gland forming’ regions transcriptionally corresponded to the ‘basal-like’ PDAC subtype, with significantly poorer outcomes ^13^.

Cancer-associated fibroblasts (CAFs) and other cell types of the tumour microenvironment have additionally emerged as key contributors to PDAC development and therapy resistance ^14,15^. In recent single-cell transcriptomics studies, at least three distinct subtypes of CAFs have been described in human and murine tumour samples: immunosuppressive cytokine-secreting inflammatory CAFs (iCAFs), myofibroblastic CAFs (myCAF) with high expression of α-smooth muscle actin *(ACTA2)* that produce extracellular matrix and are thought to restrain tumour growth, and antigen-presenting CAFs (apCAFs) expressing MHC class II and CD74 that may play an immunomodulatory role ^16–21^. Distinctive stromal gene expression signatures as well as patterns of immune cell and vascular infiltration have also been described ^4,22,23^. How these different microenvironment cell types interact with PDAC tumour cells remains a key area of research; their complex cellular exchanges may comprise both tumour-enhancing and tumour-suppressive effects, and curtail or boost the efficacy of therapeutic approaches ^24,25^.

Investigating the relationship between tumour morphology, transcriptionally different PDAC subtype cells and their microenvironment is complicated by the spatial heterogeneity and low neoplastic cell content of PDAC tumours ^23,26^. While single-cell transcriptomics has helped to distinguish diverse cell types within tumour samples, spatial information is lost during tissue dissociation. Addressing this challenge, recently developed in situ sequencing (ISS) approaches enable transcriptional profiling at the single-cell level while retaining spatial context ^27,28^.

Here, we apply ISS to probe how gene expression in single PDAC tumour cells relates to local tissue morphology. We distinguish five transcriptional subtypes of PDAC correlating with distinct morphological patterns and microenvironment cell type compositions, and show that these spatio-transcriptional subtypes hold prognostic significance.

## Methods

### Sample acquisition

Pancreatic tissue specimens from 10 patients with pancreatic ductal adenocarcinoma (PDAC) were obtained from the Tissue Biobank of Klinikum rechts der Isar and TUM (MTBIO). Tumour content was approved by a board-certified pathologist. Informed consent was available from all patients. The use of tumour material was approved by the ethics committee of the medical faculty of TUM (403/17S).

### In situ sequencing

To spatially characterise gene expression across patients, formalin-fixed paraffin-embedded tissue sections (5 μm thick) were processed for RNA in situ hybridisation according to the manufacturer’s instructions (HS Library Prep Kit Large 1110-02, CARTANA, 10xGenomics), with four DNA probes for each target gene (with a total of 199 different target genes) designed and manufactured by CARTANA (10xGenomics). As a modification, to enhance the probe signal to background ratio, 1× Lipofuscin Autofluorescence Quencher (Promocell) was applied for 30 seconds prior to fluorescence labelling. Sequencing was performed by CARTANA (10xGenomics) in six subsequent rounds of fluorescent labelling and stripping to detect the spatial coordinates of each target probe. On average, around 420,000 transcripts per sample (~190,000 – ~905,000) passed high threshold quality control. A reference 4’,6-diamidino-2-phenylindole (DAPI) staining image was also acquired for each sample. Finally, all slides were stained using an adapted hematoxylin and eosin (H&E) staining protocol ^29^.

### Analysis

#### ISS data pre-processing

Given the DAPI stained images for each sample, the nuclei were detected and segmented using the deep learning framework StarDist version 0.7.2 for object detection with star-convex polygons ^30^. The neural network was pre-trained on fluorescent nuclear marker images based on a subset of the DSB 2018 nuclei segmentation challenge dataset ^30^. Cell boundaries were approximated by isotropically expanding the nuclei labels to the maximum radius of 6 μm, with the constraint of prohibiting overlaps of cells (Supplementary Figure 1). The number of cells per sample ranged from 71,666 to 270,454. Transcripts detected with ISS were assigned to these cells by mapping the coordinates of the sequenced target probes to the cell boundaries, resulting in a cell × gene count matrix. Online visualisations were generated using the Python package TissUUmaps version 3.0.9^31^.

#### Transcriptomic analysis

ISS data were processed using the Python package Squidpy version 1.1.2 ^32^. Cells with less than four detected transcripts and transcripts detected in less than ten cells were excluded from the analysis. Transcript counts per cell were normalised and log-transformed and scaled to unit variance and zero mean. To identify cell types, a reduced set of transcripts representing cell type markers was used (Supplementary Table 3). PCA was performed on the transcript counts of cells from all patient samples and a neighbourhood graph was constructed based on the first 20 principal components. Clusters were identified by Leiden clustering (resolution = 2.0). The cellular identity of clusters was then determined based on differentially expressed genes. Clusters corresponding to the same cell type based on marker gene expression were merged, while clusters comprising two distinct cell types were split by subclustering. Transcriptional profiles were visualised using UMAP ^33^ for dimensional reduction. For more detailed analysis, clustering was repeated for malignant PDAC cells, fibroblasts, immune cells, endocrine cells and exocrine cells separately. A small number of cells which could not be identified as any specific cell type were excluded from further analysis (5.1% of all cells). PCA and clustering were also performed for cells from each patient sample individually to confirm that the combined clustering was representative.

#### Spatial co-occurrence analysis

To analyse enrichment and depletion of cell types as a function of distance from other cell types, a graph encoding spatial neighbour relations was constructed, including neighbours within a distance of 50 μm. An enrichment score was calculated based on the connectivity graph by comparing the number of observed cell type co-occurrences against 1,000 random permutations and computing a z-score. Enrichment z-scores were visualised as heatmaps for each patient sample.

#### Spatial correlation analysis

To measure spatial co-occurrence of cell types, spatial auto-correlation and cross-correlation analysis was performed using the R package MERINGUE version 1.0 ^34^. For co-occurrence at the level of transcripts, a hexagonal grid spaced at 100 μm distance between hexagon centres was defined spanning each sample, and transcripts were assigned to the nearest grid point. Transcripts detected at less than ten grid points were excluded. Counts per grid point were normalised with a scale factor of 6,000. A binary adjacency weight matrix was computed for each sample considering grid points up to 200 μm apart as neighbours. To detect spatially correlated transcripts in each sample separately, Moran’s I as a measure of spatial cross-correlation was calculated for all neighbouring pairs and genes were summarised into spatial patterns across the population of *N* cells using the spatial cross-correlation index (SCI) as defined in MERINGUE,

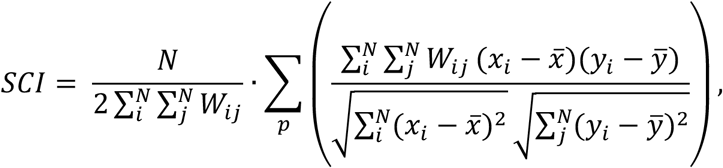

where *x* and *y* correspond to the expression magnitude of two genes in a given cell *i* and its spatially adjacent neighbours *j*. To identify spatial cross-correlation patterns across all ten samples, a joint adjacency weight matrix *W^joint^* was defined where *W^joint_i,j_^* = 1 if the indices *i* and *j* correspond to neighbouring grid points from the sample sample, and *W^joint_i,j_^* = 0 otherwise. A joint spatial cross-correlation index *SCI^joint^* was then calculated for each gene pair as

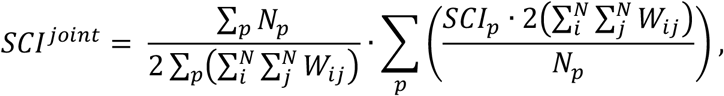

where *p* indexes samples. After calculating *SCI* and *SCI^joint^* for each gene pair, the resulting spatial cross-correlation matrices were used to group genes into spatial patterns by hierarchical clustering with Ward’s clustering criterion and dynamic tree cutting (tuning parameter deepSplit = 2).

#### Survival analysis

To analyse how identified PDAC subtype signatures relate to clinical outcomes, bulk RNA sequencing data from the TCGA-PAAD dataset ^26^ was downloaded using the R package TCGA2STAT version 1.2 ^35^ and the PACA-AU dataset (release 28) ^36^ was downloaded from the ICGC data portal (https://dcc.icgc.org/). Only samples characterised as primary solid tumour were considered (n=178 for TCGA-PAAD and n=80 for PACA-AU). RPKM expression data were converted to TPM, scaled, centred, and clipped at [-5, 5].

Expression scores for pattern-defined gene sets and ‘basal-like’ subtype marker genes (Supplementary Table 4) were determined by calculating the average expression level of each gene set for each sample, subtracted by the average expression of a control gene set. To control for differential overall expression levels of genes, all genes were binned based on average expression across all samples into 24 bins, and the control gene set was assembled by randomly selecting 100 genes from the same expression bin for each gene in the query gene set.

For each gene set, samples were divided into high and low expression groups using the median expression score as the cutoff. Kaplan-Meier plots were generated using the survminer package version 0.4.6 in R. To assess survival differences and hazard ratios, the log-rank test and cox proportional hazards regression model were used as implemented in the R package survival version 3.1-8.

## Results

### Identification of PDAC cells and pancreatic cell types using ISS

To investigate the relationship between transcriptional profiles and spatial architecture of the different cell types present in PDAC tumours, we obtained surgically resected tumour samples from ten randomly selected patients with a confirmed diagnosis of PDAC and processed them for ISS with a curated list of 199 target transcripts (Figure 1A; Supplementary Table 1 and 2). Briefly, during ISS, transcript-specific padlock probes hybridise directly to the mRNA target and are amplified and sequenced using fluorophore conjugated detection probes ^27,28,37^. Transcripts were assigned to cells by mapping their coordinates to the cell boundary map generated based on nuclei segmentation via star-convex polygons using a convolutional neural network ^30^ (Supplementary Figure 1A).

**Figure 1:**
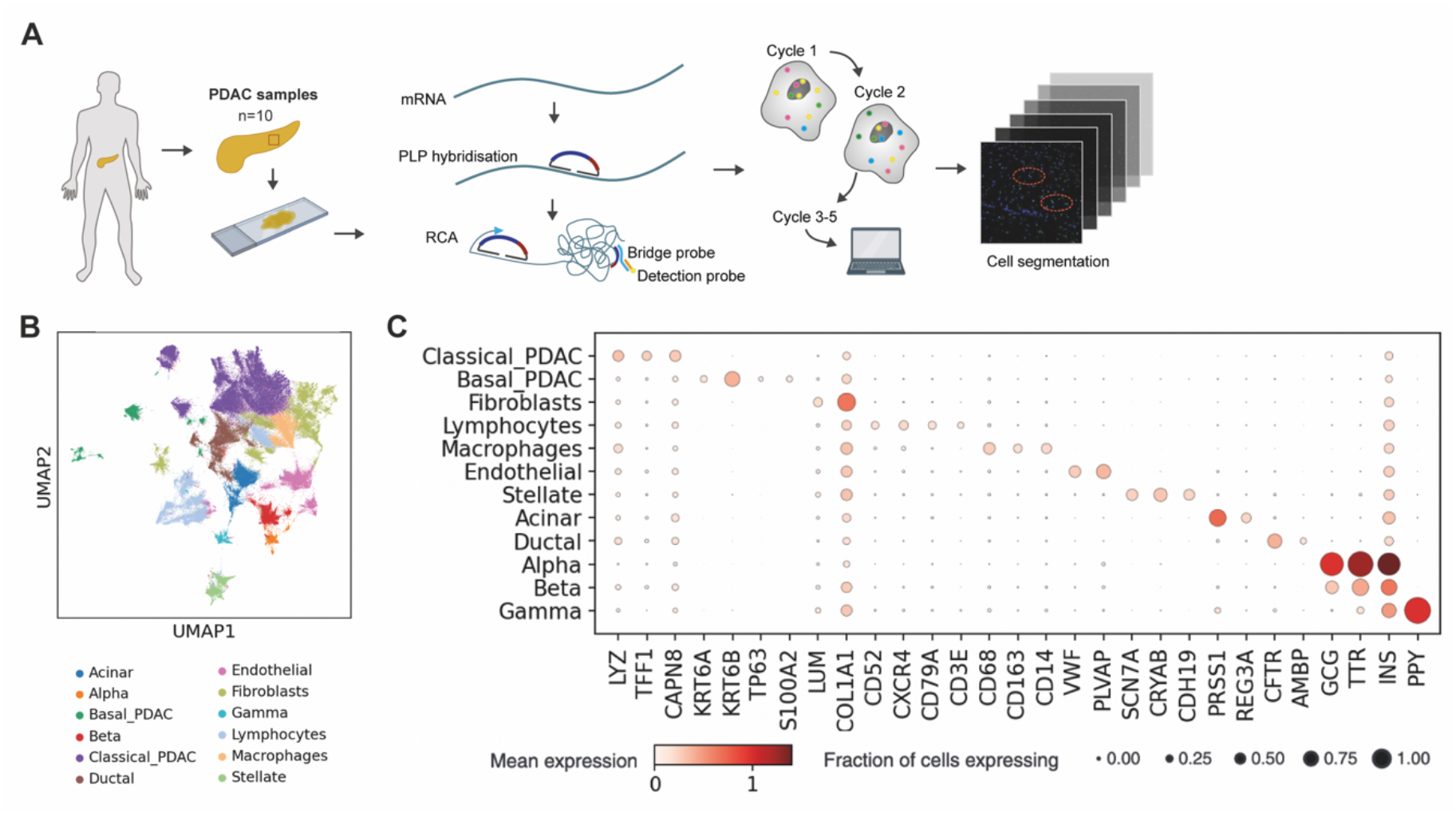
Cell type identification based on ISS of human PDAC. **(A)** Overview of the experimental workflow. PDAC biopsies were acquired from 10 patients. For in situ sequencing, tissue sections are prepared from formalin-fixed paraffin-embedded samples. Transcript-specific padlock probes (PLP) hybridise directly to the mRNA targets and are amplified by Rolling Circle Amplification (RCA). PLP identities are then decoded by sequential cycles of hybridisation and stripping of bridge and fluorophore conjugated detection probes ^27,28^. To assign transcripts to cells, a convolutional neural network is applied that segments cell nuclei via star-convex polygons ^30^. **(B)** UMAP representation of all profiled cells from ten patient samples, indicating the assigned cell types based on clustering and marker gene expression analysis. **(C)** Expression of characteristic genes across the different cell types identified in the tumour samples. Colour indicates normalised mean expression while dot size represents the fraction of cells in each population expressing the gene.

By unsupervised clustering of the resulting single-cell transcriptional profiles, we identified different cell types present in the samples based on differential expression of characteristic marker genes across all patients (Figure 1B,C). PDAC cells were distinguished from healthy ductal cells by elevated keratin 19 (*KRT19*) expression; using published subtype marker gene sets ^4^, clusters of PDAC cells were further classified as ‘basal-like’ or ‘classical’ based on expression of 11 ‘basal-like’ and 19 ‘classical’ marker genes (Supplementary Table 2). Consistent with previous reports ^8^, the neoplastic cell content of samples was relatively low (41% across all patients).

In addition to PDAC tumour cells, we identified a prominent fibroblast compartment based on expression of lumican (*LUM*), collagens and other fibroblast marker genes, which comprised multiple subclusters. Reflecting immune infiltration of the tumour volumes, lymphocytes and macrophages were also detected. Endothelial cells were classified based on von Willebrand factor (*VWF*) expression and Schwann cells based on expression of sodium channel protein type 7 subunit alpha (*SCN7A*) as well as crystallin alpha B (*CRYAB*). All samples also contained endocrine pancreatic islet cells which could be further subdivided into alpha cells expressing glucagon (*GCG*) and transthyretin (*TTR*), beta cells marked by insulin (*INS*) expression and absence of the other endocrine markers, gamma cells expressing pancreatic polypeptide (*PPY*), and a small number of delta cells marked by expression of somatostatin (*SST*). Finally, exocrine pancreatic acinar cells were identified by serine protease 1 (*PRSS1*), amylase alpha 2A (*AMY2A*) and regenerating family member 3 alpha (*REG3A*) expression.

The different cell types were represented in broadly similar proportions across patients (Supplementary Figure 1B).

By recording spatial coordinates for each detected transcript, ISS enables the morphological characterisation of cells in their spatial environment within the tissue. As expected, endocrine cells were found to occur in localised clumps representing pancreatic islets, confirming the validity of our experimental approach and processing pipeline (Supplementary Figure 1C).

### Spatial architectures of PDAC subtype cells

PDAC tumour cells from the ten patient samples did not present as a homogeneous cell population, but separated into distinct clusters based on gene expression (Figure 1B). To differentiate PDAC cell subpopulations, we performed unsupervised clustering of the malignant cells alone. Besides ‘basal-like’ PDAC cells, four clusters of PDAC cells corresponding to the ‘classical’ subtype were distinguished based on differential gene expression (Figure 2A,B and Supplementary Figure 2A).

**Figure 2:**
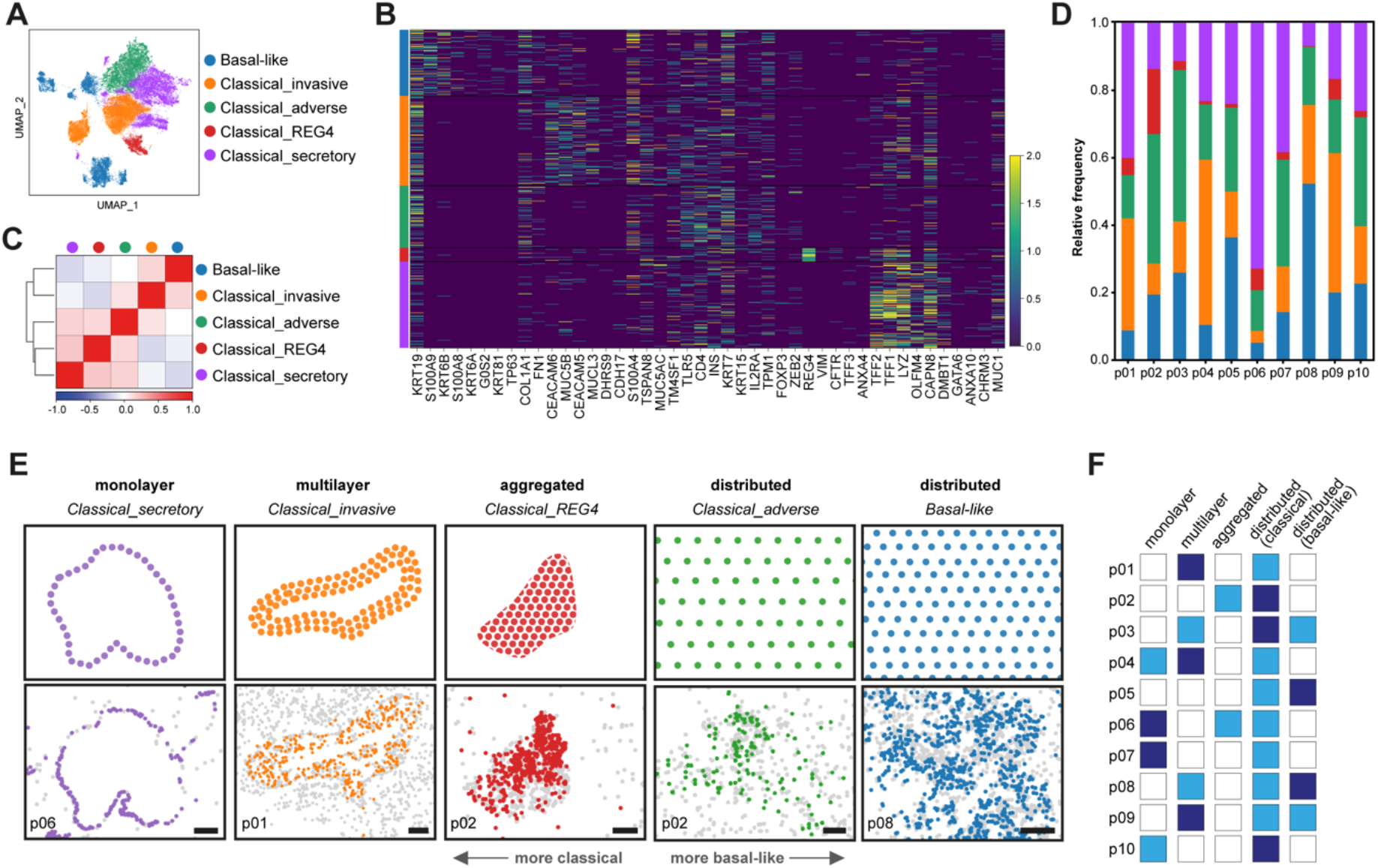
Transcriptional subtypes of PDAC with characteristic morphologies. **(A)** UMAP representation of the identified PDAC subtypes, labelled according to functional annotations of enriched genes in each cluster (see also Supplementary Figure 2A). **(B)** Pearson’s correlation coefficients between mean transcriptional profiles of the different PDAC subtypes. **(C)** Normalised expression of characteristic genes, including marker genes for the ‘classical’ and ‘basal-like’ subtypes, across the identified PDAC cell clusters. Colour bars denote subtypes, with colours as in (A). **(D)** Relative frequencies of the different PDAC subtypes across all ten patient samples. Colours indicate subtypes as in (A). **(E)** Distinctive morphologies of the identified transcriptional PDAC subtypes were observed across patients. Top row depicts simplified illustrations of morphologies, while bottom row shows representative areas from different samples where the respective PDAC subtype and morphology was detected. Colours indicate PDAC subtypes as in (A), with all other tumour cells shown in grey. Scale bars, 200 μm. **(F)** Representation of morphological PDAC subtypes across patient samples. Dark blue: dominant morphology in the sample, light blue: morphology also detected in the sample, white: morphology not detected.

One major cluster was characterised by the expression of secretion related genes including *LYZ, TFF1* and *TFF2*, and was therefore labelled *Classical_secretory*. Another large cluster comprised cells with high expression of the carcinoembryonic antigen-related cell adhesion molecules *CEACAM5* and *CEACAM6*, which mediate cell adhesion and promote tumour invasion ^38^, as well as *MUCL3*, which has been shown to enhance PDAC cell proliferation, migration and invasion ^39^; this was labelled *Classical_invasive*. A smaller cluster defined by high expression of *KRT7*, an intermediate filament protein known to be overexpressed in pancreatic cancer tissues compared to non-malignant pancreatic tissue ^40^, was labelled *Classical_adverse* since a correlation of *KRT7* overexpression with poorer overall survival of PDAC patients has been suggested ^41^. Finally, the smallest cluster comprised cells highly expressing regenerating family member 4 (*REG4*), which is thought to play a role in carcinoma development from intestinal-type intraductal papillary mucinous neoplasms (IPMNs) ^42^. The role of *REG4* in PDAC progression remains unclear; serum *REG4* levels in PDAC patients have been shown to predict unfavourable histologic response to neoadjuvant chemoradiotherapy and a higher rate of postoperative local recurrence ^43^, but a recent study also associated *REG4* expression with longer survival ^44^. This cluster was labelled *Classical_REG4*.

Transcriptional correlation confirmed that these clusters represent distinct PDAC cell states, with *Classical_adverse* showing greater transcriptional similarity with ‘basal-like’ PDAC compared to the other ‘classical’ PDAC cell states (Figure 2C). While none of the identified PDAC cell states derived from a single patient of origin, they were differently represented across patients (Figure 2D). ‘Basal-like’ PDAC cells were mostly detected in patient samples 05 and 08 (36% and 52% of all PDAC cells in these samples, respectively). Among ‘classical’ PDAC subclusters, *Classical_REG4* cells were most prominent in sample 02 (19%), *Classical_secretory* cells in samples 01 and 06 (40% and 73%) and *Classical_invasive* cells in samples 04 and 09 (49% and 41%).

The spatial information retained in ISS data enables the assessment of the local architecture of these distinct PDAC cell states (Supplementary Figure 2B). Remarkably, across the ten-patient cohort, we observed recurring characteristic differences in tumour morphology related to the dominant PDAC cell state in different slide regions (Figure 2E,F and Supplementary Figure 3). ‘Basal-like’ PDAC cells, which mostly derived from two patients, were diffusely distributed across contiguous areas of the tumour tissue. ‘Classical’ tumour cells were detected either as monolayers around a lumen, multiple layers of cells around a lumen, clumps, or distributed across tissue regions. *Classical_REG4* cells largely presented in clumps, which could also comprise *Classical_secretory* cells. *Classical_secretory* cells were otherwise mostly detected as monolayers around a lumen, histologically representing more highly differentiated tumour areas, unless they co-localised with *Classical_invasive* cells and shared the spatial architecture of the latter. *Classical_invasive* cells presented as multiple layers of cells around a lumen where one existed, or distributed across areas without a lumen. *Classical_adverse* cells, on the other hand, which were detected in all patient samples, exhibited a distributed morphological pattern similar to ‘basal-like’ PDAC cells.

Notably, different PDAC cell states co-occurred within individual patients, but with a phenotypic gradient from ‘classical’ to ‘basal-like’ states. *Classical_secretory* cells, characterised by secretion related gene expression and their monolayer morphology closely resembling healthy pancreatic ducts, did not co-occur with ‘basal-like’ PDAC cells. In contrast, *Classical_invasive* cells and ‘basal-like’ PDAC cells were observed in different regions of the same tumour section in samples 03, 08 and 09. As *Classical_invasive* cells showed increased gene expression associated with adhesion and invasion, this suggests a continuum of transcriptional and morphological states, with *Classical_secretory* and ‘basal-like’ PDAC cells occupying the opposite ends while *Classical_invasive* cells correspond to an intermediate phenotype. Upregulation of *CEACAM5, CEACAM6* and *MUCL3* may disrupt an initially formed PDAC cell monolayer, leading to the emergence of multilayer structures and dissemination of tumour cells throughout the surrounding tissue.

### Co-localisation of malignant and tumour microenvironment cells

PDAC development and prognosis is intricately linked with the tumour microenvironment, including cancer-associated fibroblasts (CAFs), infiltrating immune cells and vasculature ^15^. Unsupervised clustering of fibroblasts from all patient samples revealed four distinct clusters (Figure 3A,B). Inflammatory CAFs (iCAFs) were identified based on expression of hyaluronan synthase 1 (*HAS1*) and interleukin 6 (*IL6*), while myofibroblastic CAFs (CAFs) subdivided into two clusters. One cluster was distinguished by periostin (*POSTN*) expression, encoding an integrin ligand that supports cell adhesion and migration ^45^; it was labelled *myCAF_adhesive*. Enrichment for α-smooth muscle actin (*ACTA2*) expression characterised the second myCAF cluster, which was therefore labelled *myCAF_contractile*.

**Figure 3:**
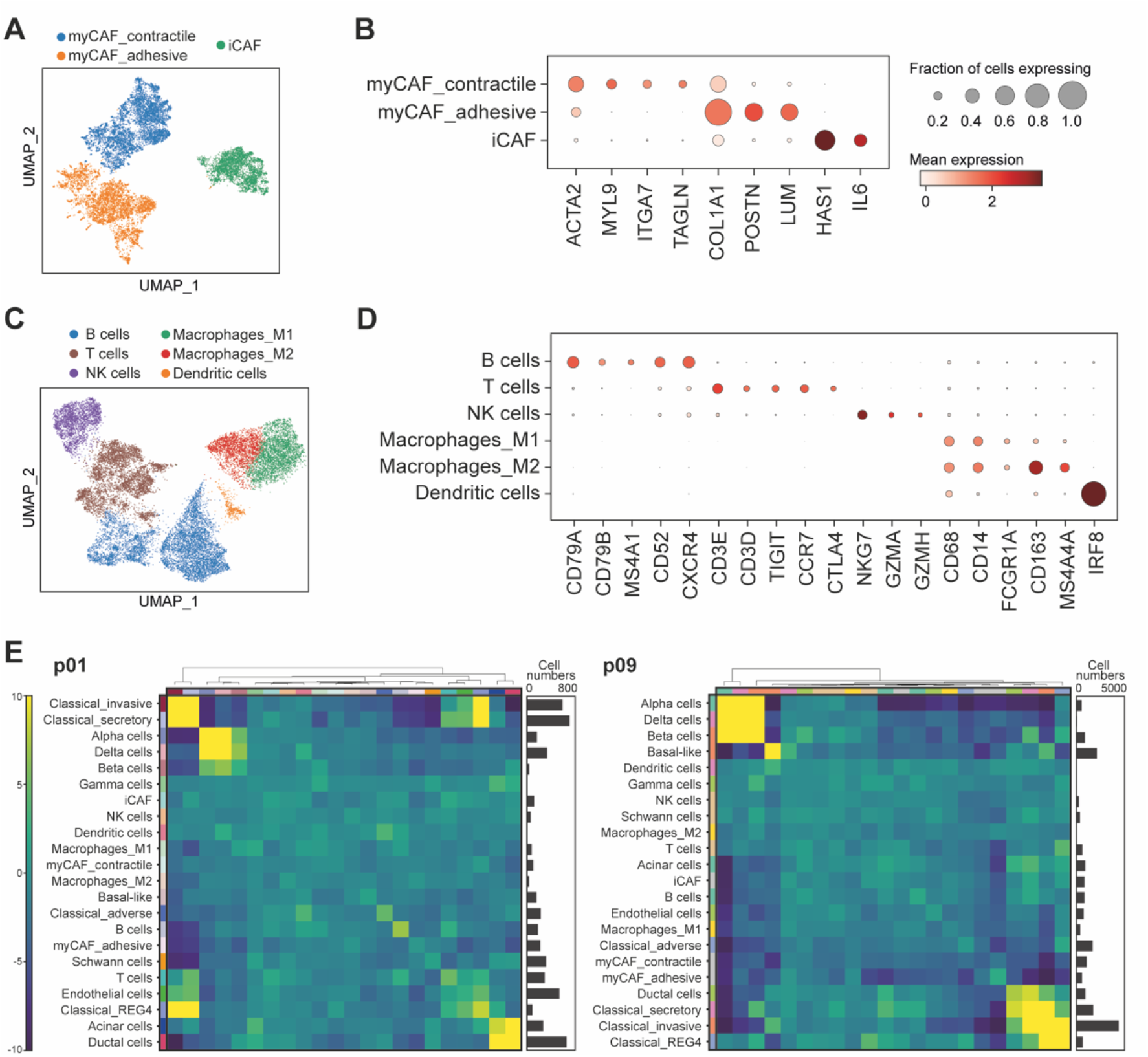
Stromal and immune cell types in PDAC tumour samples. **(A)** UMAP representation of CAF populations, including inflammatory CAFs as well as two types of myofibroblastic CAFs. **(B)** Expression of characteristic genes across the CAF populations. **(C)** UMAP representation of immune cell populations, including macrophages, B cells, T cells and NK cells. **(D)** Expression of characteristic genes across the immune cell populations. **(E)** Enrichment or depletion of cell types in local neighbourhoods was assessed by comparing the number of observed cell type co-occurrences within 50 μm against expected values based on random permutations on the cell connectivity graph. Heatmaps show z-scores for enrichment or depletion of cell type pairings for two patient samples. Bar plots indicate cell numbers for each sample.

While the limited number of transcripts in ISS experiments did not allow the distinction of immune cell subtypes at high resolution, the major categories of immune cells were also identified (Figure 3C,D). Macrophages, marked by expression of *CD68* and *CD14*, could be divided into pro-inflammatory M1 macrophages expressing *FCGR1A* and regulatory M2 macrophages with increased expression of *CD163* and *MS4A4A*. Lymphocytes comprised large clusters of B cells and T cells. Finally, a cluster of cells enriched for expression of cytotoxicity-related genes including granzymes (*GZMA, GZMH*) and *NKG7* was identified as NK cells, although it cannot be ruled out that it might comprise cytotoxic T cells.

To gain insight into the spatial co-localisation of PDAC tumour with microenvironment cells, we constructed a cell connectivity graph for each sample where all cells within a 50 μm radius of each other were counted as neighbours. We then quantified the enrichment of cell types in local neighbourhoods by determining the frequency of neighbouring cell type pairings and comparing it to expectation based on a randomly permuted graph ^32^. Across samples, *myCAF_adhesive* cells are largely absent from tumour areas with a clearly ‘classical’ phenotype, i.e. around *Classical_secretory* cells (Figure 3E); this is consistent with previous reports of poorer prognosis in tumours with a dominant *POSTN* expressing fibroblast population ^45,46^. Inflammatory CAF are often spatially associated with ‘basal-like’ PDAC and immune cells. Consistent with the proposed continuum of PDAC subtypes showing more ‘classical’ to more ‘basal-like’ features, the vicinity of *Classical_adverse* tumour cells is also enriched for ‘basal-like’ PDAC cells as well as immune cells (Figure 3E).

### Spatial patterns of gene expression in PDAC tumour samples

As the tumour samples in our cohort comprised varying proportions of the different PDAC subtypes, we adapted a published approach ^34^ to analyse the spatial cross-correlation of tumour cells with their microenvironment across all patients (Figure 4A). Due to the relatively low number of transcripts detected per cell, we defined a hexagonal grid with a distance of 100 μm between spots and assigned transcripts the nearest grid spots. Transcripts with spatially heterogeneous expression were identified through Local Indicators of Spatial Association (LISA) ^34,47^ using normalised transcript counts and a binary adjacency weight matrix encoding adjacent spots within a spatial distance of 200 μm. Hierarchical clustering of transcripts based on their spatial cross-correlation across all samples revealed patterns of transcripts with spatially coherent expression profiles (Supplementary Figure 4A,B).

**Figure 4:**
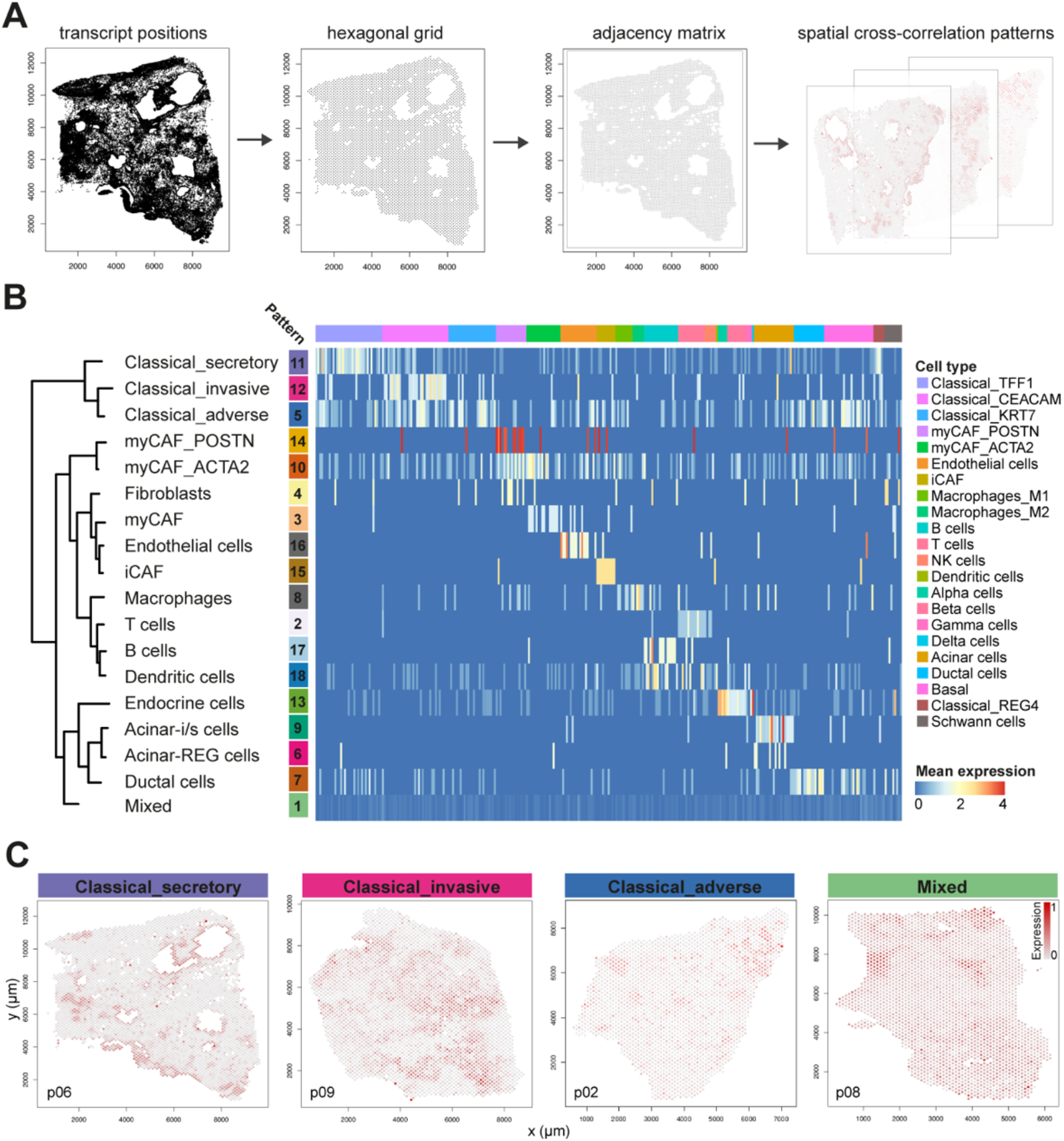
Spatial cross-correlation patterns of transcripts. **(A)** To identify spatial patterns of transcription, each recorded transcript is assigned to the nearest point on a hexagonal grid (grid point distance 100 μm). A binary adjacency weight matrix is determined for each sample, with two points considered adjacent if they are located within 200 μm of each other. Here, grey lines connect adjacent spots. Using the adjacency weight matrices for each sample, a spatial cross-correlation index is computed for every transcript pair taking into account neighbourhood information from all samples. Gene expression patterns are determined by dynamic tree cutting of a hierarchical dendrogram computed from the resulting spatial cross-correlation matrix. Groups of genes are z-scored and averaged for visualisation of transcriptional patterns ^34^. **(B)** Hierarchical dendrogram depicting patterns of spatially cross-correlated transcripts across all samples. Most patterns could be identified as representing specific cell types and were labelled accordingly; the pattern labelled ‘Mixed’ comprised *REG4*, markers for ‘basal-like’ PDAC, Schwann cells, proliferation and angiogenesis, as well as various immune signalling genes. The full dendrogram including all transcript names is shown in Supplementary Figure 4A. Heatmap shows average gene expression per pattern across all cells, with column annotations indicating cell type identity and patient origin. **(C)** Visualisation in different patient samples of spatial transcriptional patterns that correspond to the ‘classical’ PDAC subtypes, as well as the distributed pattern comprising ‘basal-like’ PDAC marker genes. Colour indicates normalised mean expression of pattern transcripts.

Spatial patterns largely reflect the previously identified cell types present in the PDAC samples (Figure 4B,C). Transcripts with high expression in endocrine cells, as expected, show high spatial cross-correlation with each other. They are spatially associated with transcripts enriched in the exocrine compartment, i.e. acinar-i/s cells^29^, acinar-REG cells^48^ and ductal cells, reflecting healthy pancreatic tissue areas (Figure 4B). Transcripts characteristic of T cells, B cells, macrophages and immune-related surface genes also co-localise, and are spatially associated with fibroblast and endothelial cell enriched transcripts. Despite their morphological differences, transcripts identifying the ‘classical’ PDAC subpopulations *Classical_secretory, Classical_invasive* and *Classical_adverse* show overlapping spatial expression profiles. In contrast, *Classical_REG4* and ‘basal-like’ PDAC transcripts do not form a separate pattern but co-localise with other transcripts showing a spatially distributed expression profile, including transcripts associated with stromal cells, Schwann cells, proliferation, angiogenesis and immune signalling (Figure 4B). This suggests that *Classical_REG4* and ‘basal-like’ PDAC populations cannot be delineated based on spatial cross-correlation analysis of the marker genes employed to identify these cell populations in our data, and additional marker genes will be required to spatially resolve these populations in future studies. Spatial cross-correlation patterns of gene expression in PDAC tumour samples thus corroborate the distinction of ‘classical’ PDAC subtypes, confirming that transcriptional differences relate to distinct morphologies.

Among fibroblasts, myCAFs show a closer spatial correlation with ‘classical’ PDAC subtypes compared to iCAF, which in turn are associated with other microenvironment cell types as well as ‘basal-like’ PDAC (Figure 4B, Figure 3E). These results indicate that presence of different PDAC tumour subpopulations induces compositional changes of the microenvironment and/or microenvironment composition affects PDAC subtype identity.

In addition to established cell type marker genes, our ISS probe set included additional targets based on their reported or postulated role in PDAC development (Supplementary Table 2). While the majority of these targets showed no coherent patterns of co-localisation with PDAC subtype specific transcripts (Supplementary Figure 4A), we observed spatial clustering of *Classical_invasive* cells with complement decay-accelerating factor (*CD55*), a glycoprotein that accelerates the decay of complement cascade proteins and thereby prevents damage to cells ^49^. In PDAC and other cancers, including colorectal and head and neck cancers, the evasion of complement cells achieved by elevated *CD55* has been shown to confer worse prognosis ^50–52^. Moreover, hypoxia-inducible factor 1 (*HIF1A*) expression was spatially associated with iCAF, in line with a recent study suggesting that hypoxia drives iCAF formation in PDAC ^53^. Finally, we found a spatial co-localisation of neuropilin-2 (*NRP2*) with myCAF. Neuropilin-2 is known as a receptor for angiogenic growth factors. Besides its expression by endothelial cells ^54^, *NRP2* in PDAC cells is associated with angiogenesis, tumour growth, migration and invasion ^55^. PDAC cells have also been reported to induce *NRP2* expression in tumour-associated macrophages, in turn promoting tumour growth ^56^. In gastric cancer, *NRP2* is upregulated in CAFs compared to normal fibroblasts and high expression levels correlate with worse outcomes ^57^; our in situ sequencing data suggest this may also be true for PDAC.

### Prognostic significance of ‘classical’ PDAC subpopulations

While it is well established that ‘basal-like’ PDAC carry a worse prognosis compared to ‘classical’ PDAC tumours, the proposed multiplicity of ‘classical’ PDAC subtypes raises the question whether phenotypic features of these subtypes could be harnessed for prognostic substratification. We addressed this question by means of the pancreatic adenocarcinoma cohort within The Cancer Genome Atlas (TCGA-PAAD), limiting our analysis to primary tumour samples (n=178) ^8^. To probe our findings in a separate cohort, we also analysed pancreatic adenocarcinoma data from the Pan-Cancer Analysis of Whole Genomes (PCAWG) study (PACA-AU), again considering only primary tumour samples (n=80) ^36^. For each sample, gene set expression scores were computed based on bulk RNA-seq data for the gene sets corresponding to *Classical_secretory*, *Classical_invasive* and *Classical_adverse* spatial patterns as well as ‘basal-like’ marker genes (Figure 5A-C and Supplementary Table 4).

**Figure 5:**
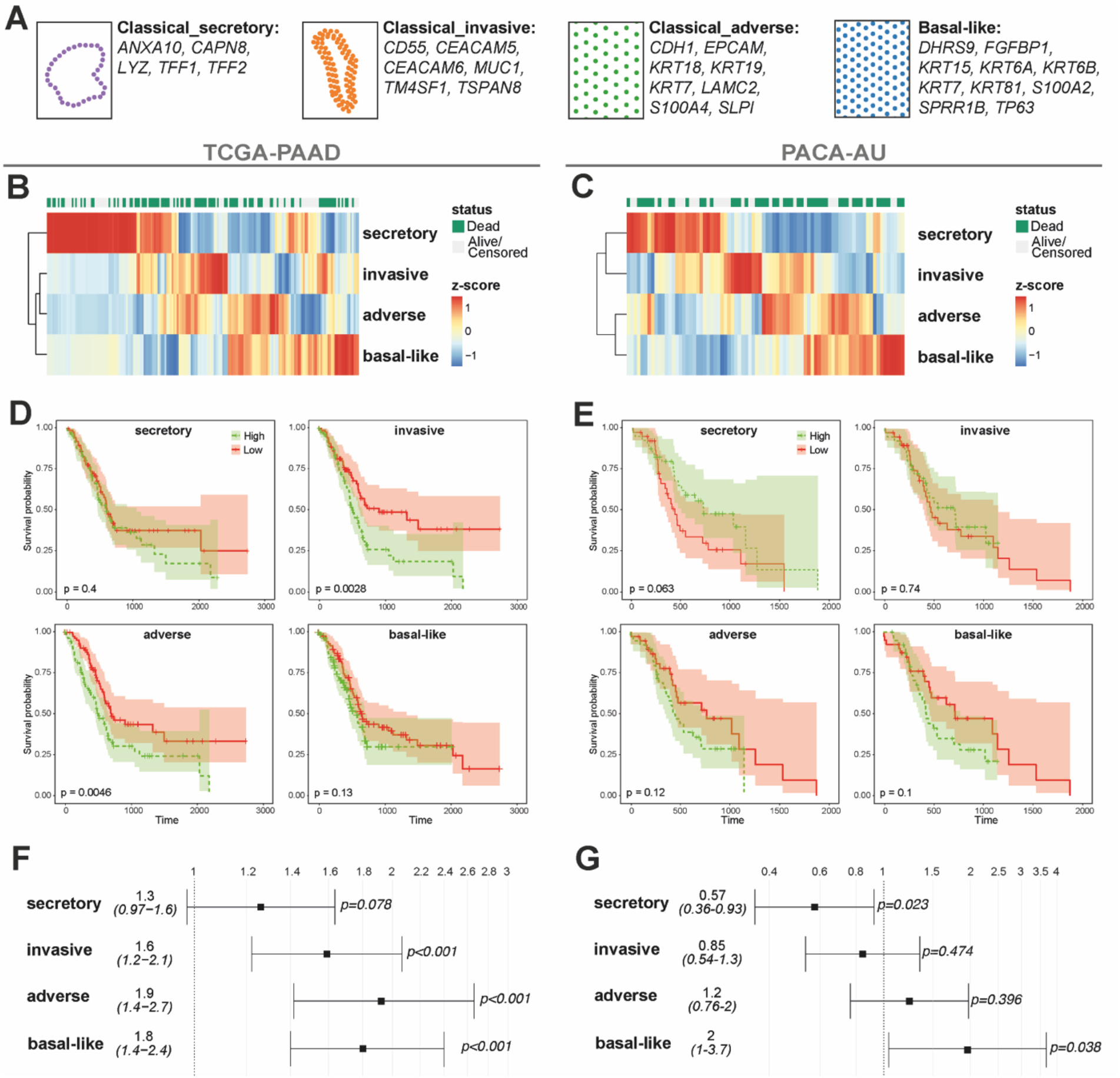
Prognostic relevance of PDAC subtypes. **(A)** Gene sets corresponding to *Classical_secretory*, *Classical_invasive* and *Classical_adverse* spatial patterns as well as ‘basal-like’ marker genes. **(B,C)** PDAC subtype scores based on bulk RNA-seq data for primary tumour samples from the TCGA-PAAD cohort (n=178 patients) ^8^ and the PACA-AU cohort (n=80 patients) ^36^, visualised by z-scores for each sample. **(D,E)** Kaplan-Meier curves comparing survival probability in TCGA-PAAD and PACA-AU patients with high (green) or low (red) expression of the gene sets defined by the spatial patterns representing secretory, invasive and adverse ‘classical’ PDAC subtypes. The comparison according to ‘basal-like’ signature gene expression is also shown for reference. Shaded areas indicate 95% confidence intervals. High scores for the invasive or adverse phenotypes are associated with significantly worse survival compared to the secretory phenotype (p<0.01, log-rank test). **(F,G)** Hazard ratios associated with gene set scores for the secretory, invasive, adverse and ‘basal-like’ gene sets. Expression of the invasive or adverse gene sets is associated with worse outcome compared to the secretory gene set (p<0.001, Wald test).

In the TCGA-PAAD cohort, no significant difference in survival was observed between patients with high or low expression of *Classical_secretory* genes, consistent with the notion that this presents the most ‘classical’ phenotype reminiscent of healthy pancreatic duct tissue (Figure 5D). In contrast, high expression of *Classical_invasive* and *Classical_adverse* genes was associated with significantly worse survival. As expected, expression of ‘basal-like’ subtype marker genes was also associated with poor outcome. Hazard ratio (HR) analysis confirmed worse outcomes associated with the *Classical_invasive* (HR 1.6, 95% confidence interval 1.2-2.1), *Classical_adverse* (HR 1.9, 95% confidence interval 1.4-2.7) and ‘basal-like’ (HR 1.8, 95% confidence interval 1.4-2.4) phenotypes (Figure 5F).

In the smaller PACA-AU cohort, survival differences were less significant, but we observed comparable tendencies (Figure 5E). Hazard ratios also indicated poorer prognosis for the ‘basal-like’ PDAC subtype (HR 2, 95% confidence interval 1-3.7) while the *Classical_secretory* subtype emerged as protective (HR 0.57, 95% confidence interval 0.36-0.93) (Figure 5G).

Overall, these results corroborate a gradient of worsening overall survival from *Classical_secretory* to *Classical_invasive* and *Classical_adverse* tumours. As *Classical_secretory* tumour cells exhibit transcriptional features of healthy pancreatic tissue while *Classical_adverse* cells are most transcriptionally similar to ‘basal-like’ PDAC cells, we conclude that consistent survival differences, associated with transcriptional subtypes, exist even within configurations traditionally referred to as ‘classical’ PDAC.

## Discussion

Intratumoural heterogeneity remains a significant obstacle to PDAC treatment. In this study, we employed in situ sequencing to identify subpopulations of PDAC with distinct transcriptional and morphological characteristics. Our findings suggest a further stratification of ‘classical’ PDAC, which represent the majority of PDAC cases ^7^, into four subtypes representing a continuum from more ‘classical’ to more ‘basal-like’ phenotypes. Morphologically, we observed a spatial association of ‘classical’ PDAC subtypes, whereas ‘basal-like’ PDAC are distributed in the stroma and co-localise preferentially with iCAF; this is consistent with the distinction between ‘classical’ and ‘squamoid-basaloid’ spatial communities in a recent whole-transcriptome profiling study ^58^.

Among the ‘classical’ PDAC cell populations, *Classical_secretory* cells most closely resemble healthy pancreatic ductal tissue, both in terms of morphology and gene expression. *Classical_REG4* cells are distinguished by high expression of REG4 but otherwise transcriptionally similar to *Classical_secretory* cells; while the latter largely occur as monolayers around a lumen, *Classical_REG4* cells present as cell aggregates within the samples. *REG4* has been suggested as a potential serological marker of PDAC and a target for antibody therapy ^59^. Our data suggests that *REG4* overexpression may be limited to a subset of PDAC cells, potentially restricting the utility of this approach. Interestingly, *REG4* has been linked to PDAC development from intestinal-type intraductal papillary mucinous neoplasms (IPMNs) ^42^, a potential alternative cancerogenic route that might be reflected in the different morphologies of *Classical_secretory* and Classical_REG4 tumour cell populations.

*Classical_invasive* cells are characterised by increased expression of carcinoembryonic antigen-related cell adhesion molecules *CEACAM5* and *CEACAM6* along with multi-layer or more distributed tumour architectures. *CEACAM5* and *CEACAM6* expression reportedly correlates with shortened overall and disease-free survival in PDAC, as well as positive lymph node status and distant metastasis ^60,61^. Their expression is also associated with the progression of pancreatic intraepithelial neoplasia (PanIN), its most common precursor lesion, to malignant PDAC ^38^. Moreover, *CEACAM6* has been linked to the invasive capacity of PDAC cells *in vitro* ^62,63^. Together with the described PDAC subtype morphologies, this suggests that *CEACAM5* and *CEACAM6* expression triggers the capacity of tumour cells to part from ductal monolayer structures and invade into the surrounding tissue or disseminate to distant sites. Interestingly, in an immunohistochemistry study of PDAC tissue microarrays, *CEACAM5* and *CEACAM6* expression was higher in moderately-differentiated than in well-differentiated or poorly-differentiated tumours ^64^, potentially reflecting the intermediate state that *Classical_invasive* cells occupy between more ‘classical’ and more ‘basal-like’ PDAC subtypes. Finally, *Classical_adverse* cells are most ‘basal-like’ and their expression profile is enriched for a combination of ‘basal-like’ and ‘classical’ marker genes.

By stratifying the TCGA-PAAD and PACA-AU cohorts according to the morpho-transcriptional PDAC subtypes identified here, we found that overall survival decreased on a gradient from more ‘classical’ to more ‘basal-like’ tumours. Despite the small cohort size, we also observed a tendency for better survival associated with lumina in the tumours and worse survival with ‘basal-like’ tumours within our own dataset of ten patients (Supplementary Table 1). Morpho-transcriptional PDAC subtypes thus carry prognostic significance.

Our results contribute to resolving the current multitude of partially overlapping classification schemes for PDAC tumours ^4–7^ by taking into account their spatial context. Transcriptionally defined subtypes with characteristic morphological features occupy a continuum from ‘classical’ to ‘basal-like’ PDAC and co-occur within the same tumours, consistent with previous observations of tumour subtype co-existence by RNA sequencing of dissociated cells ^9,10,65^. Notably, *Classical_secretory* cells morphologically represent glandular or duct-like differentiation patterns, corresponding to higher differentiated tumour areas and better outcomes according to WHO grading criteria for PDAC, whereas *Classical_invasive* and *Classical_adverse* tumours with their invasive or distributed spatial architectures reflect morphological criteria for higher-grade tumours ^66^.

While in situ sequencing is not yet feasible for clinical applications, the correspondence between transcriptional and morphological features of PDAC might in future enable the automated substratification of PDAC tumours based on morphology alone, for example using stained tumour sections acquired as part of routine clinical procedures. In addition, more comprehensive profiling using whole-transcriptome spatial analysis at the single-cell level could uncover molecular interactions between the different PDAC subtypes and their microenvironment, aiding the development of targeted therapies for PDAC.

## Acknowledgements

The survival analysis presented here is in part based upon data generated by the TCGA Research Network (https://www.cancer.gov/tcga) and the International Cancer Genome Consortium (https://dcc.icgc.org/); we gratefully acknowledge the respective clinical contributors and data producers. We thank the Tissue Biobank of MRI and TUM as well as Anja Kühl and Simone Spieckermann at iPATH.Berlin for excellent technical support, and Naveed Ishaque and Sebastian Tiesmeyer for helpful discussions.

## Author contributions

CC, KS, WW and RE conceived of and supervised the project. FS, BK KS and LT prepared PDAC samples. LT, MR, XQ and AN designed probes and conducted experiments. TGK, AS and JL analysed data. TGK wrote the manuscript with input from AS and FS. WW, KS, RE and CC acquired funding. All authors commented on the manuscript.

## Supplementary Figures

**Supplementary Figure 1:**
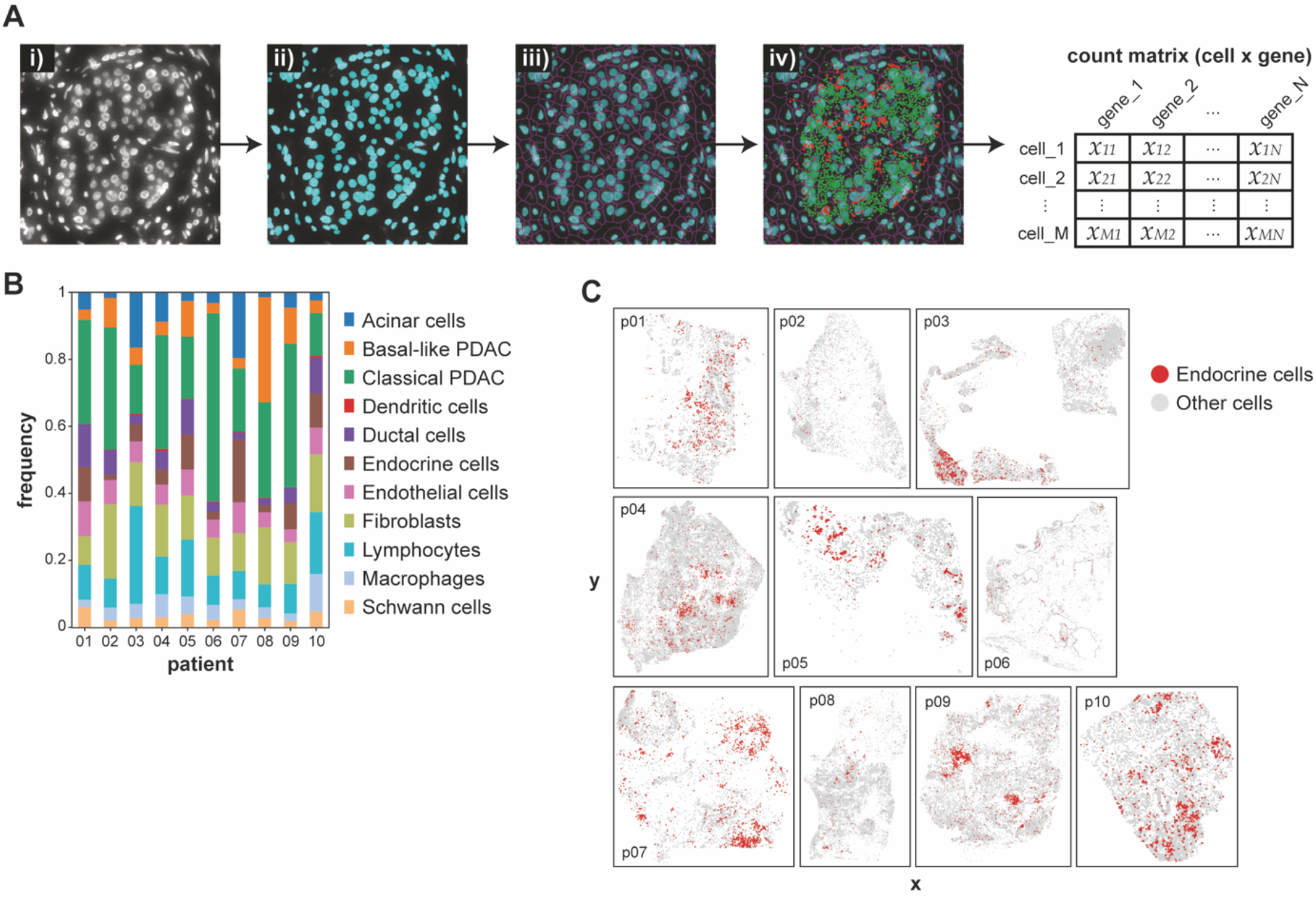
Characterisation of cell types detected in PDAC samples. **(A)** Assignment of ISS transcript locations to cells using the deep-learning based segmentation framework StarDist: The DAPI stained nuclei image (i) is segmented using StarDist (ii). The nuclei label mask is expanded isotropically without overlapping to approximate cell boundaries (iii) and the transcript locations are mapped onto the cell boundary map (iv), resulting in a cell × gene count matrix. **(B)** Frequency of cell types detected across all patient samples. **(C)** Spatial locations of endocrine cells reflecting pancreatic islets and other detected cells across all patient samples.

**Supplementary Figure 2:**
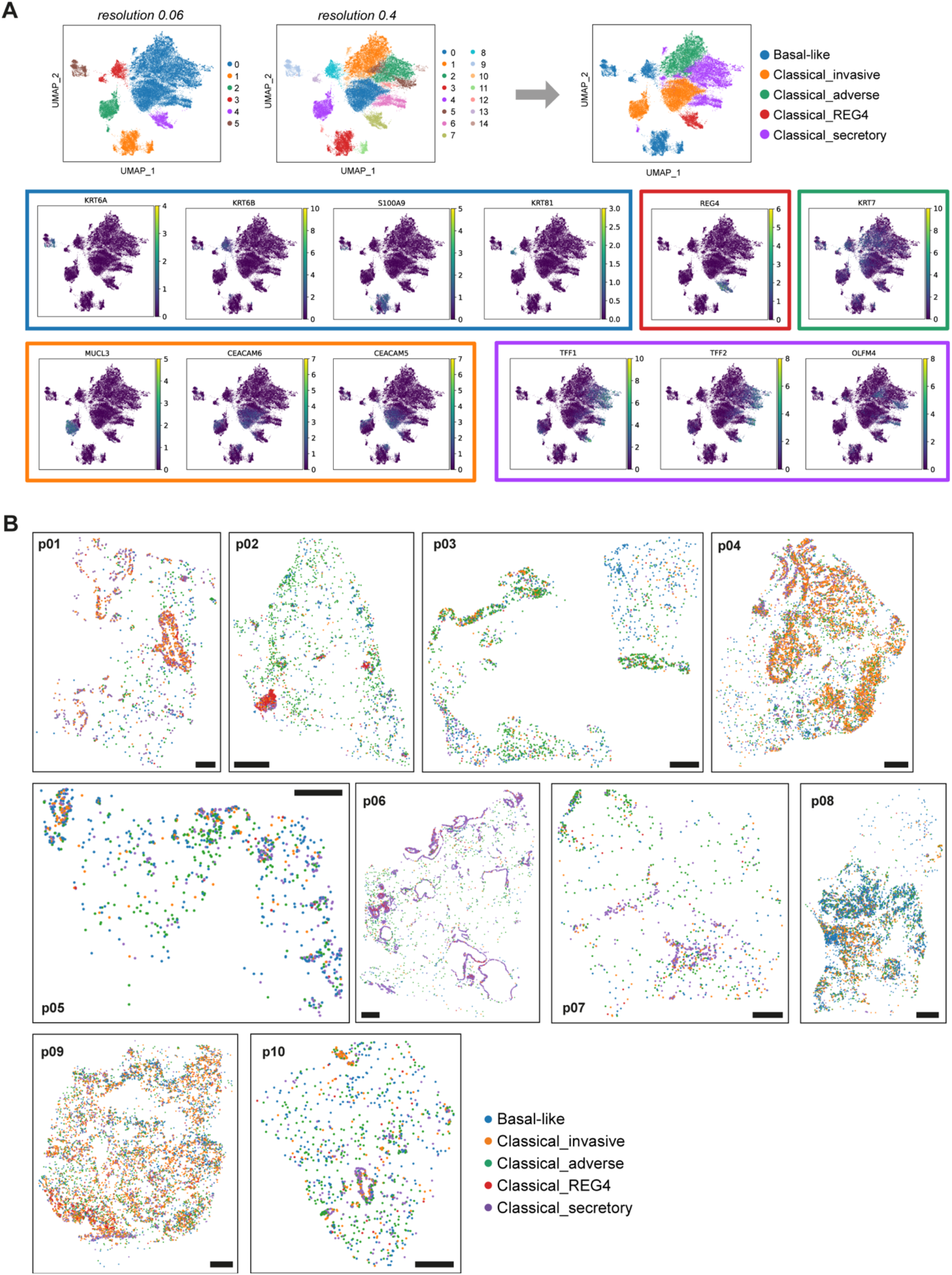
Identification of PDAC subtypes and their spatial distribution. **(A)** PDAC subtype identity was determined based on Leiden clustering and marker gene expression. At lower clustering resolution (0.06), clusters 0, 2 and 4 expressed ‘classical’ marker genes but were distinguished by enrichment for *MUCL3* and *CEACAM6* in cluster 2 and *REG4* in cluster 4, while the remaining clusters expressed ‘basal-like’ marker genes. At higher clustering resolution (0.4), fifteen clusters were identified by Leiden clustering. Of these, clusters 3, 8, 9, 11, 13, were enriched for ‘basal-like’ marker gene expression and were therefore summarised as ‘Basal-like’. Among the ‘classical’ clusters, clusters 0 and 4 shared enrichment for invasion related genes such as *CEACAM5* and *CEACAM6* and were summarised as *Classical_invasive;* clusters 2, 5, 6, 10, 12, 14, shared expression of secretion related genes such as *TFF1, TFF2* and *OLFM4* and were therefore summarised as *Classical_secretory*; cluster 1 did not show secretory features but high expression of *KRT7*, associated with poorer outcome in PDAC ^41^, and was therefore labelled *Classical_adverse;* and the REG4-expressing cluster 7 was labelled *Classical_REG4*. Cell clusters and gene expression are visualised on the same UMAP representation as in Figure 2A. Coloured boxes correspond to the identified PDAC subtypes. **(B)** Spatial distribution of PDAC subtype cells across patient samples. Each dot represents a PDAC tumour cell coloured by subtype identity, with the spatial distribution corresponding to the ISS coordinates. Scale bars, 1 mm.

**Supplementary Figure 3:**
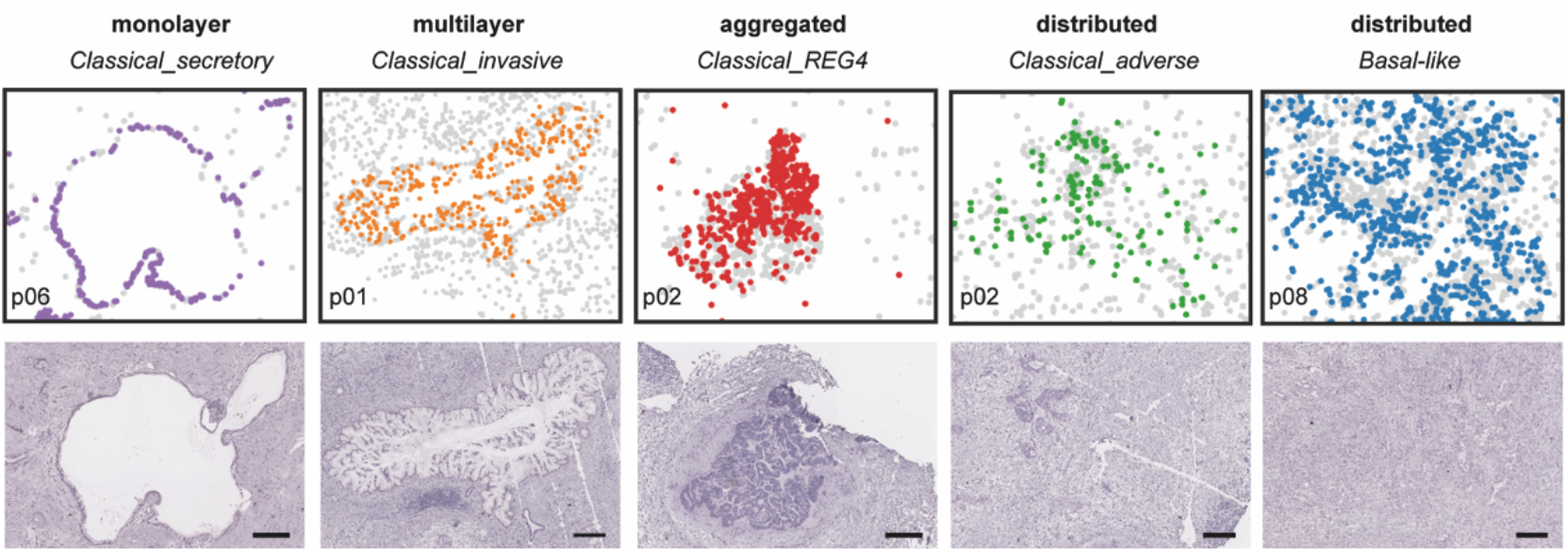
Identified PDAC morphologies as shown in Figure 2E (top) and H&E staining images of the corresponding tumour areas (bottom). Scale bars, 200 μm.

**Supplementary Figure 4:**
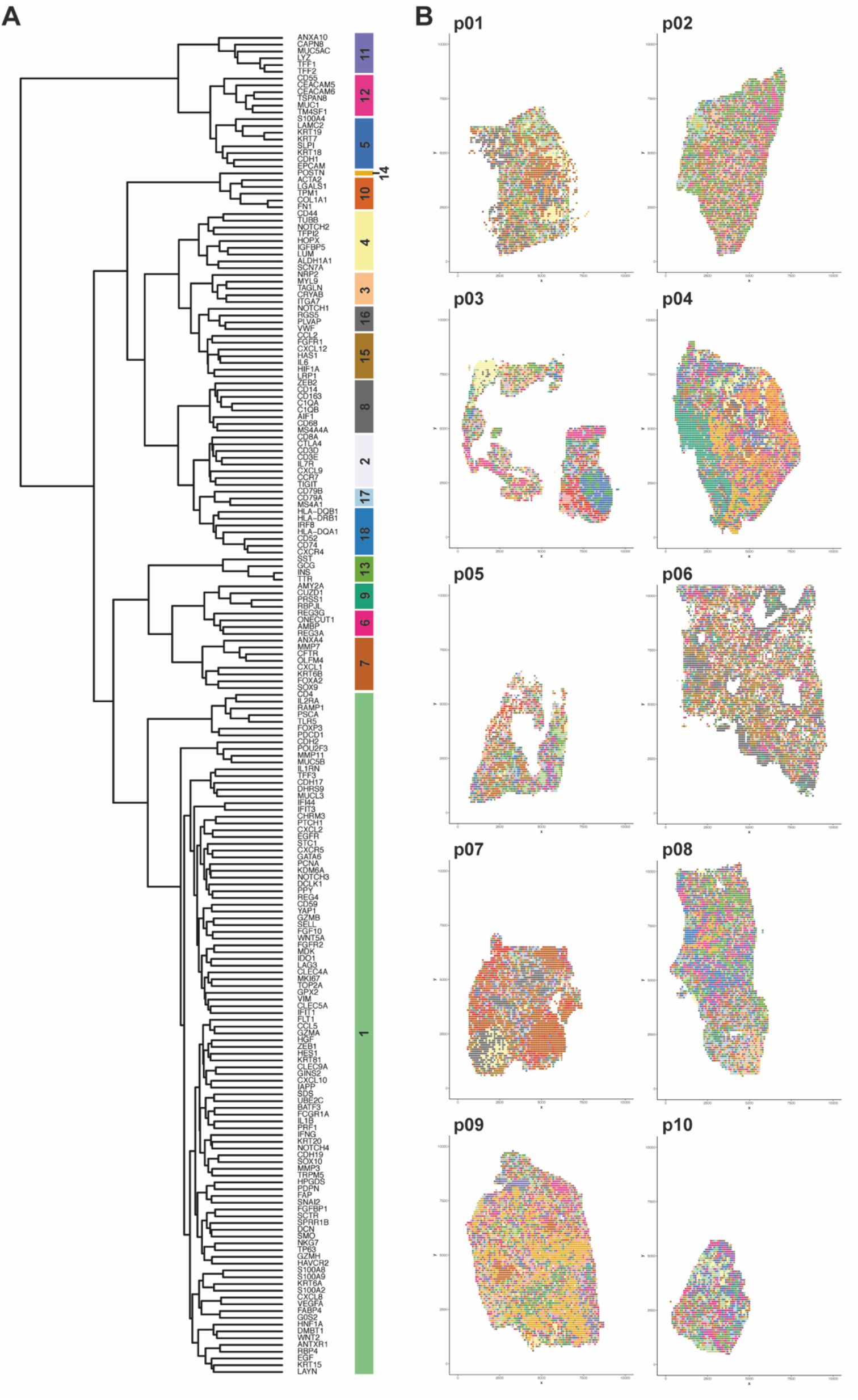
**(A)** Hierarchical clustering for all transcripts based on spatial cross-correlation across all samples. **(B)** Distribution of dominant patterns across samples. Each position is coloured according to the locally dominant pattern.

## References

1. Bengtsson, A., Andersson, R. & Ansari, D. The actual 5-year survivors of pancreatic ductal adenocarcinoma based on real-world data. Sci. Rep. 10, 1–9 (2020).

2. Siegel, R. L., Miller, K. D., Fuchs, H. E. & Jemal, A. Cancer statistics, 2022. CA. Cancer J. Clin. 72, 7–33 (2022).

3. Rahib, L. et al. Projecting cancer incidence and deaths to 2030: The unexpected burden of thyroid, liver, and pancreas cancers in the united states. Cancer Res. 74, 2913–2921 (2014).

4. Moffitt, R. A. et al. Virtual microdissection identifies distinct tumor-and stroma-specific subtypes of pancreatic ductal adenocarcinoma. Nat. Genet. 47, 1168–1178 (2015).

5. Collisson, E. A. et al. Subtypes of pancreatic ductal adenocarcinoma and their differing responses to therapy. Nat. Med. 17, 500–503 (2011).

6. Bailey, P. et al. Genomic analyses identify molecular subtypes of pancreatic cancer. Nature 531, 47–52 (2016).

7. Collisson, E. A., Bailey, P., Chang, D. K. & Biankin, A. V. Molecular subtypes of pancreatic cancer. Nat. Rev. Gastroenterol. Hepatol. 16, 207–220 (2019).

8. Raphael, B. J. et al. Integrated Genomic Characterization of Pancreatic Ductal Adenocarcinoma. Cancer Cell 32, 185–203.e13 (2017).

9. Juiz, N. et al. Basal-like and classical cells coexist in pancreatic cancer revealed by single-cell analysis on biopsy-derived pancreatic cancer organoids from the classical subtype. FASEB J. 34, 12214–12228 (2020).

10. Krieger, T. G. et al. Single-cell analysis of patient-derived PDAC organoids reveals cell state heterogeneity and a conserved developmental hierarchy. Nat. Commun. 12, (2021).

11. Chan-seng-yue, M. et al. Transcription phenotypes of pancreatic cancer are driven by genomic events during tumor evolution. Nat. Genet. doi:10.1038/s41588-019-0566-9.

12. Lokuhetty, D., White, V. A., Watanabe, R. & Cree, I. A. WHO Classification of Tumours: Digestive System Tumours. (International Agency for Research on Cancer, 2019).

13. Kalimuthu, S. N. et al. Morphological classification of pancreatic ductal adenocarcinoma that predicts molecular subtypes and correlates with clinical outcome. Gut 69, 317–328 (2020).

14. Biffi, G. & Tuveson, D. A. Diversity and biology of cancer-associated fibroblasts. Physiol. Rev. 101, 147–176 (2021).

15. Ho, W. J., Jaffee, E. M. & Zheng, L. The tumour microenvironment in pancreatic cancer — clinical challenges and opportunities. Nat. Rev. Clin. Oncol. 17, 527–540 (2020).

16. Öhlund, D. et al. Distinct populations of inflammatory fibroblasts and myofibroblasts in pancreatic cancer. J. Exp. Med. 214, 579–596 (2017).

17. Elyada, E. et al. Cross-Species Single-Cell Analysis of Pancreatic Ductal Adenocarcinoma Reveals Antigen-Presenting Cancer-Associated Fibroblasts. Cancer Discov. 9, 1102–1123 (2019).

18. Dominguez, C. X. et al. Single-cell RNA sequencing reveals stromal evolution into LRRC15+ myofibroblasts as a determinant of patient response to cancer immunotherapy. Cancer Discov. 10, 232–253 (2020).

19. Biffi, G. et al. Il1-induced Jak/STAT signaling is antagonized by TGFβ to shape CAF heterogeneity in pancreatic ductal adenocarcinoma. Cancer Discov. 9, 282–301 (2019).

20. Bernard, V. et al. Single-cell transcriptomics of pancreatic cancer precursors demonstrates epithelial and microenvironmental heterogeneity as an early event in neoplastic progression. Clin. Cancer Res. 25, 2194–2205 (2019).

21. Hosein, A. N. et al. Cellular heterogeneity during mouse pancreatic ductal adenocarcinoma progression at single-cell resolution. JCI Insight 4, (2019).

22. Maurer, C. et al. Experimental microdissection enables functional harmonisation of pancreatic cancer subtypes. Gut 68, 1034–1043 (2019).

23. Vickman, R. E. et al. Deconstructing tumor heterogeneity: The stromal perspective. Oncotarget 11, 3621–3632 (2020).

24. Klemm, F. & Joyce, J. A. Microenvironmental regulation of therapeutic response in cancer. Trends Cell Biol. 25, 198–213 (2015).

25. Grünwald, B. T. et al. Spatially confined sub-tumor microenvironments in pancreatic cancer. Cell 184, 5577–5592.e18 (2021).

26. Raphael, B. J. et al. Integrated Genomic Characterization of Pancreatic Ductal Adenocarcinoma. Cancer Cell 32, 185–203.e13 (2017).

27. Ke, R. et al. In situ sequencing for RNA analysis in preserved tissue and cells. Nat. Methods 10, 857–860 (2013).

28. Gyllborg, D. et al. Hybridization-based in situ sequencing (HybISS) for spatially resolved transcriptomics in human and mouse brain tissue. Nucleic Acids Res. 48, E112 (2020).

29. Tosti, L. et al. Single-Nucleus and In Situ RNA–Sequencing Reveal Cell Topographies in the Human Pancreas. Gastroenterology 160, 1330–1344.e11 (2021).

30. Schmidt, U., Weigert, M., Broaddus, C. & Myers, G. Cell detection with star-convex polygons. Lect. Notes Comput. Sci. (including Subser. Lect. Notes Artif. Intell. Lect. Notes Bioinformatics) 11071 LNCS, 265–273 (2018).

31. Solorzano, L., Partel, G. & Wählby, C. TissUUmaps: Interactive visualization of large-scale spatial gene expression and tissue morphology data. Bioinformatics 36, 4363–4365 (2020).

32. Palla, G. et al. Squidpy: a scalable framework for spatial omics analysis. Nat. Methods 19, 171–178 (2022).

33. McInnes, L., Healy, J. & Melville, J. UMAP: Uniform Manifold Approximation and Projection for Dimension Reduction. arXiv 1802.03426v2 (2018).

34. Miller, B. F., Bambah-Mukku, D., Dulac, C., Zhuang, X. & Fan, J. Characterizing spatial gene expression heterogeneity in spatially resolved single-cell transcriptomic data with nonuniform cellular densities. Genome Res. 31, 1843–1855 (2021).

35. Wan, Y. W., Allen, G. I. & Liu, Z. TCGA2STAT: Simple TCGA data access for integrated statistical analysis in R. Bioinformatics 32, 952–954 (2016).

36. Scarlett, C. J., Salisbury, E. L., Biankin, A. V. & Kench, J. Precursor lesions in pancreatic cancer: Morphological and molecular pathology. Pathology 43, 183–200 (2011).

37. Lee, H., Marco Salas, S., Gyllborg, D. & Nilsson, M. Direct RNA targeted in situ sequencing for transcriptomic profiling in tissue. Sci. Rep. 12, 1–9 (2022).

38. Zińczuk, J. et al. Expression of chosen carcinoembryonic-related cell adhesion molecules in pancreatic intraepithelial neoplasia (PanIN) associated with chronic pancreatitis and pancreatic ductal adenocarcinoma (PDAC). Int. J. Med. Sci. 16, 583–592 (2019).

39. Yan, J. et al. High expression of diffuse panbronchiolitis critical region 1 gene promotes cell proliferation, migration and invasion in pancreatic ductal adenocarcinoma. Biochem. Biophys. Res. Commun. 495, 1908–1914 (2018).

40. Bournet, B. et al. Gene expression signature of advanced pancreatic ductal adenocarcinoma using low density array on endoscopic ultrasound-guided fine needle aspiration samples. Pancreatology 12, 27–34 (2012).

41. Li, Y., Su, Z., Wei, B. & Liang, Z. Krt7 overexpression is associated with poor prognosis and immune cell infiltration in patients with pancreatic adenocarcinoma. Int. J. Gen. Med. 14, 2677–2694 (2021).

42. Nakata, K. et al. REG4 is associated with carcinogenesis in the ‘intestinal’ pathway of intraductal papillary mucinous neoplasms. Mod. Pathol. 22, 460–468 (2009).

43. Eguchi, H. et al. Serum REG4 level is a predictive biomarker for the response to preoperative chemoradiotherapy in patients with pancreatic cancer. Pancreas 38, 791–798 (2009).

44. Bhardwaj, A. et al. Deeper insights into long-term survival heterogeneity of Pancreatic Ductal Adenocarcinoma (PDAC) patients using integrative individual-and group-level transcriptome network analyses. bioRxiv 2020.06.01.116194 (2020) doi:10.1101/2020.06.01.116194.

45. Liu, Y. et al. Role of microenvironmental periostin in pancreatic cancer progression. Oncotarget 8, 89552–89565 (2017).

46. Neuzillet, C. et al. Inter-and intra-tumoural heterogeneity in cancer-associated fibroblasts of human pancreatic ductal adenocarcinoma. J. Pathol. 248, 51–65 (2019).

47. Anselin, L. Local Indicators of Spatial Association—LISA. Geogr. Anal. 27, 93–115 (1995).

48. Muraro, M. J. et al. A Single-Cell Transcriptome Atlas of the Human Pancreas. Cell Syst. 3, 385–394.e3 (2016).

49. Spendlove, I., Ramage, J. M., Bradley, R., Harris, C. & Durrant, L. G. Complement decay accelerating factor (DAF)/CD55 in cancer. Cancer Immunol. Immunother. 55, 987–995 (2006).

50. He, Z., Wu, H., Jiao, Y. & Zheng, J. Expression and prognostic value of CD97 and its ligand CD55 in pancreatic cancer. Oncol. Lett. 9, 793–797 (2015).

51. Durrant, L. G. et al. Enhanced expression of the complement regulatory protein CD55 predicts a poor prognosis in colorectal cancer patients. Cancer Immunol. Immunother. 52, 638–642 (2003).

52. Kesselring, R. et al. The complement receptors CD46, CD55 and CD59 are regulated by the tumour microenvironment of head and neck cancer to facilitate escape of complement attack. Eur. J. Cancer 50, 2152–2161 (2014).

53. Mello, A. et al. Hypoxia promotes an inflammatory phenotype of fibroblasts in pancreatic cancer. bioRxiv (2022).

54. Islam, R. et al. Role of Neuropilin - 2 - mediated signaling axis in cancer progression and therapy resistance. Cancer Metastasis Rev. (2022) doi:10.1007/s10555-022-10048-0.

55. Dallas, N. A. et al. Neuropilin-2-mediated tumor growth and angiogenesis in pancreatic adenocarcinoma. Clin. Cancer Res. 14, 8052–8060 (2008).

56. Roy, S. et al. Macrophage-derived neuropilin-2 exhibits novel tumor-promoting functions. Cancer Res. 78, 5600–5617 (2018).

57. Yang, Y. et al. CAF promotes chemoresistance through NRP2 in gastric cancer. Gastric Cancer 25, 503–514 (2022).

58. Hwang, W. L. et al. Single-nucleus and spatial transcriptome profiling of pancreatic cancer identifies multicellular dynamics associated with neoadjuvant treatment. Nat. Genet. 54, (2022).

59. Takehara, A. et al. Novel tumor marker REG4 detected in serum of patients with resectable pancreatic cancer and feasibility for antibody therapy targeting REG4. Cancer Sci. 97, 1191–1197 (2006).

60. Gebauer, F. et al. Carcinoembryonic antigen-related cell adhesion molecules (CEACAM) 1, 5 and 6 as biomarkers in pancreatic cancer. PLoS One 9, (2014).

61. Duxbury, M. S., Ito, H., Zinner, M. J., Ashley, S. W. & Whang, E. E. CEACAM6 gene silencing impairs anoikis resistance and in vivo metastatic ability of pancreatic adenocarcinoma cells. Oncogene 23, 465–473 (2004).

62. Okuda, R. et al. Reconstructing cell interactions and state trajectories in pancreatic cancer stromal tumoroids. bioRxiv 2022.02.14.480334 (2022).

63. Chen, J. et al. CEACAM6 induces epithelial-mesenchymal transition and mediates invasion and metastasis in pancreatic cancer. Int. J. Oncol. 43, 877–885 (2013).

64. Blumenthal, R. D., Leon, E., Hansen, H. J. & Goldenberg, D. M. Expression patterns of CEACAM5 and CEACAM6 in primary and metastatic cancers. BMC Cancer 7, 8809–8817 (2007).

65. Topham, J. T. et al. Subtype-discordant pancreatic ductal adenocarcinoma tumors show intermediate clinical and molecular characteristics. Clin. Cancer Res. 27, 150–157 (2021).

66. Nagtegaal, I. D. et al. The 2019 WHO classification of tumours of the digestive system. Histopathology 76, 182–188 (2020).

